# Expanding the toolbox for hypervirulent *Klebsiella pneumoniae* using the social amoeba *Dictyostelium discoideum* as a virulence biosensor

**DOI:** 10.1101/2025.11.14.688115

**Authors:** Ian Perez, Matías Gálvez-Silva, Diego Rojas, Antonia Ramos, Miranda Bravo, Yahua Chen, Yunn-Hwen Gan, Francisco P. Chávez, Andrés E. Marcoleta

## Abstract

Hypervirulent *Klebsiella pneumoniae* (hvKp) represents a critical global threat due to its capacity to cause severe, community-acquired infections in healthy individuals and its convergence with multidrug resistance. However, current methods to evaluate hvKp virulence are limited by their reliance on mammalian models, which are costly, ethically constrained, and unsuitable for high-throughput applications. Here, we introduce a biosensing approach using the social amoeba *Dictyostelium discoideum* as a living phenotypic sensor for hvKp virulence. Through a suite of quantitative assays integrating predation resistance, social development, and real-time single-cell motility tracking, we show that *D. discoideum* can discriminate avirulent, virulent, and hypervirulent *K. pneumoniae* strains. Using the model hvKp strain SGH10 and its capsule-null mutant Δ*wcaJ*, we demonstrate that capsule production is strongly associated with phagocytosis resistance, developmental arrest, and amoebae paralysis, all quantifiable through imaging and motility-based readouts. Multivariate data integration revealed a clear segregation of virulence phenotypes, validating *D. discoideum* as a sensitive, scalable, and genetically tractable biosensor for functional virulence detection. This work establishes a conceptual and methodological framework for developing cell-based biosensors that report virulence levels through measurable host behavioral responses and outlines a calibration path to vertebrate endpoints. These systems provide a low-cost alternative to mammalian systems, especially for early-stage evaluation, and a potential foundation for automated high-throughput screening and surveillance of virulence in critical priority bacterial pathogens.

## INTRODUCTION

*K. pneumoniae* is a Gram-negative enterobacterium found in different environmental matrices and in mammalian mucosal surfaces. Initially associated mainly with mild nosocomial infections in immunocompromised patients (Podschun and Ullmann, 1998), currently, multidrug-resistant strains are ranked as the uppermost priority bacterial pathogens for global health, particularly strains resistant to the last-resort carbapenem antibiotics (Sati et al., 2025; Xu et al., 2017). Moreover, hypervirulent strains (hvKp) causing severe community-acquired infections in immunocompetent patients have emerged (Choby et al., 2020; Marr and Russo, 2019), including convergent strains showing both multidrug resistance and hypervirulence (Lam et al., 2021; Wyres et al., 2020). In February 2024, the European Center for Disease Prevention and Control (ECDC) issued an alert due to the increasing number of hvKp cases associated with high morbidity and mortality (European Centre for Disease Prevention and Control, 2024), emphasizing the importance of hvKp early detection and the need to prevent their spread in healthcare settings. Among the vulnerabilities identified against this threat, especially in low-to middle-income countries, are the limited local epidemiological and genomic information of these clones, and insufficient capacity for their detection.

HvKp strains were first isolated in the 1980s from pyogenic liver abscesses in Taiwan, China, and Southeast Asia (Liu et al., 1986), observing that these strains can spread metastatically, causing meningitis, endophthalmitis, endocarditis, and necrotizing fasciitis, among other severe and often fatal infections (Russo and Marr, 2019). Molecular and genomic epidemiology studies have revealed that hvKp strains are characterized by the expression of an expanded set of virulence factors, mainly acquired by horizontal gene transfer, including a variety of siderophores, the genotoxin colibactin, and the antimicrobial peptide microcin E492, among others (Lam et al., 2018; Tan et al., 2024).

The thick *K. pneumoniae* capsule is one of the most prominent virulence factors in this species, promoting immune system evasion by hindering relevant surface antigens and conferring resistance to phagocytosis by macrophages and other immune cells (Liu et al., 2025; Paczosa and Mecsas, 2016). hvKp commonly show capsule overproduction and a hypermucoviscous phenotype, due to the activity of the proteins encoded in the *rmpADC* operon. RmpA is a transcriptional activator that upregulates the transcription of the *rmpC* and *rmpD* genes. RmpC then acts as a positive regulator for the capsular polysaccharide synthesis genes, resulting in capsule overproduction (Walker et al., 2019). The *rmpD* gene encodes a small protein that interacts with Wzc, a tyrosine kinase that regulates the polymerization and export of the capsular polysaccharide chains. By binding to Wzc, RmpD modifies its function, leading to the production of longer, more uniform CPS chains. This hypermucoviscous capsule acts as an antiadhesion and mechanical clearance resistance factor, promoting dissemination and resilience inside the host (Ovchinnikova et al., 2023; Walker et al., 2020).

Capsule-null *K. pneumoniae* mutants have been reported as attenuated or avirulent in different infection models, although controversy arose regarding possible pleiotropic effects on cell stability depending on the mutated gene (Fu et al., 2025; Paczosa and Mecsas, 2016; Wei et al., 2025). In this line, Tan et al. showed that mutants in *wcaJ*, encoding a glycosyltransferase responsible for the first polymerization step in the production of the capsule, are non-capsulated and avirulent in mice, retaining an intact cell envelope compared to other capsule-null mutants like Δ*wza* or Δ*wzy*, which exhibit severe membrane defects (Tan et al., 2020). This study was performed using *K. pneumoniae* SGH10, proposed as a model hvKp strain isolated from a human liver abscess (Lam et al., 2018). Hence, Δ*wcaJ* was proposed as a physiologically stable and interpretable hvKp attenuated model strain to be used as a negative control for host interaction studies. Nevertheless, until now, it has only been probed on mice (Tan et al., 2020).

Despite the rapid expansion of genomic and bioinformatic resources, genome sequencing alone remains insufficient to predict bacterial virulence with accuracy (Beck et al., 2025). The mere presence of virulence-associated genes, such as those encoding capsules, siderophores, or toxins, does not necessarily translate into a hypervirulent phenotype. Likewise, phenotypic assays such as hypermucoviscosity testing, while valuable, often capture only superficial traits that fail to reflect the complexity of host–pathogen interactions. As a result, only mammalian models currently allow reliable evaluation of virulence in hvKp (Russo et al., 2025, 2024). However, these models are constrained by ethical, financial, and scalability limitations, underscoring the urgent need for alternative systems capable of delivering functional virulence assessments in a quantitative and accessible manner.

Alternative infection models such as *Danio rerio* (zebrafish) have been widely validated for assessing virulence in various pathogens, including *K. pneumoniae* (Franza et al., 2024; Gálvez-Silva et al., 2025; Marcoleta et al., 2018). A simpler yet powerful model is the social amoeba *Dictyostelium discoideum*, a professional phagocyte that naturally feeds on bacteria and can be infected by virulent strains (Annesley and Fisher, 2009; Bozzaro and Eichinger, 2011). This protist transitions between unicellular and multicellular stages depending on environmental cues. In its single-celled phase, it engulfs bacteria via phagocytosis, closely mimicking the behavior of mammalian macrophages through conserved molecular mechanisms (Bozzaro and Eichinger, 2011). Beyond phagocytosis, *D. discoideum* exhibits immune-like functions such as micropinocytosis (Annesley et al., 2011), chemotaxis (Escalante and Vicente, 2000), and the formation of extracellular DNA traps (Zhang et al., 2016).

Experimentally, *D. discoideum* offers multiple advantages: it grows axenically with minimal equipment and enables virulence comparisons through its social development response to bacterial feeding and depletion. It has become a valuable model for studying development, multicellularity, cell signaling, and innate immunity (Dunn et al., 2018). Moreover, *D. discoideum* has proven to be an effective phenotypic biosensor of bacterial virulence, providing both mechanistic insights and high-throughput screening capabilities. Notable examples include a *D. discoideum*-*Mycobacterium marinum* platform for high-throughput anti-infective compound screening (Nitschke et al., 2025) and its use as a host for antivirulence assays in *P. aeruginosa* PAO1 (Bravo-Toncio et al., 2016). Collectively, these studies position *D. discoideum* as a biosensor for phenotypic virulence evaluation, offering an ethical, scalable, and mechanistically informative alternative to mammalian infection models.

The global emergence of hvKp underscores the need for expanded tools for its study and early detection. Although genomic analyses are increasingly accessible, they often fail to correlate reliably with phenotypic outcomes, highlighting the importance of simple, cost-effective *in vivo* infection models. Although the *Galleria mellonella* model has been widely used with hvKp (Bruchmann et al., 2021; Xiao et al., 2025), it offers only a single lethality readout and fails to distinguish classically virulent (cvKp) from hvKp (Russo and MacDonald, 2020). Previously, we validated *D. discoideum* as a model to compare virulence among cvKp strains (Marcoleta et al., 2018), and later demonstrated its utility in uncovering the role of polyphosphate in hvKp pathogenicity (Rojas et al., 2024). Building on these findings, we now establish *D. discoideum* as a robust and accessible host to study hvKp, developing an expanded suite of assays and readouts capable of differentiating attenuated, virulent, and hypervirulent *K. pneumoniae*, using well-characterized bacterial strains.

## RESULTS

### *In vitro* and *in vivo* validation of the capsule-null mutant *K. pneumoniae* SGH10 Δ*wcaJ* as an attenuated hvKp control

The validation of novel virulence assays for hvKp requires a proper attenuated control to compare with a thoroughly characterized wild-type strain. Therefore, we selected the hypervirulent strain *K. pneumoniae* SGH10 and its mutant derivative Δ*wcaJ*, demonstrated to be highly virulent and attenuated in murine infection models, respectively (Tan et al., 2020). We generated a scarless deletion of the *wcaJ* gene starting from the SGH10 strain (Figure S1) and compared the production of virulence factors *in vitro* in both strains. As expected, Δ*wcaJ* displayed significantly reduced capsule production, as revealed by uronic acid quantification (Figure 1A). As a control, we used the capsule-free *E. coli* BL21 strain. Moreover, low-speed sedimentation indicated a markedly reduced mucoviscosity in the mutant strain (Figure 1B), as expected for capsule-null cells. In accordance, scanning electron microscopy (SEM) showed that hypermucoviscous capsule loss correlated with notorious changes in the Δ*wcaJ* cell surface, which appears smooth, in contrast to the spiky surface of the wild type cells (Figure 1C).

**Figure 1.**
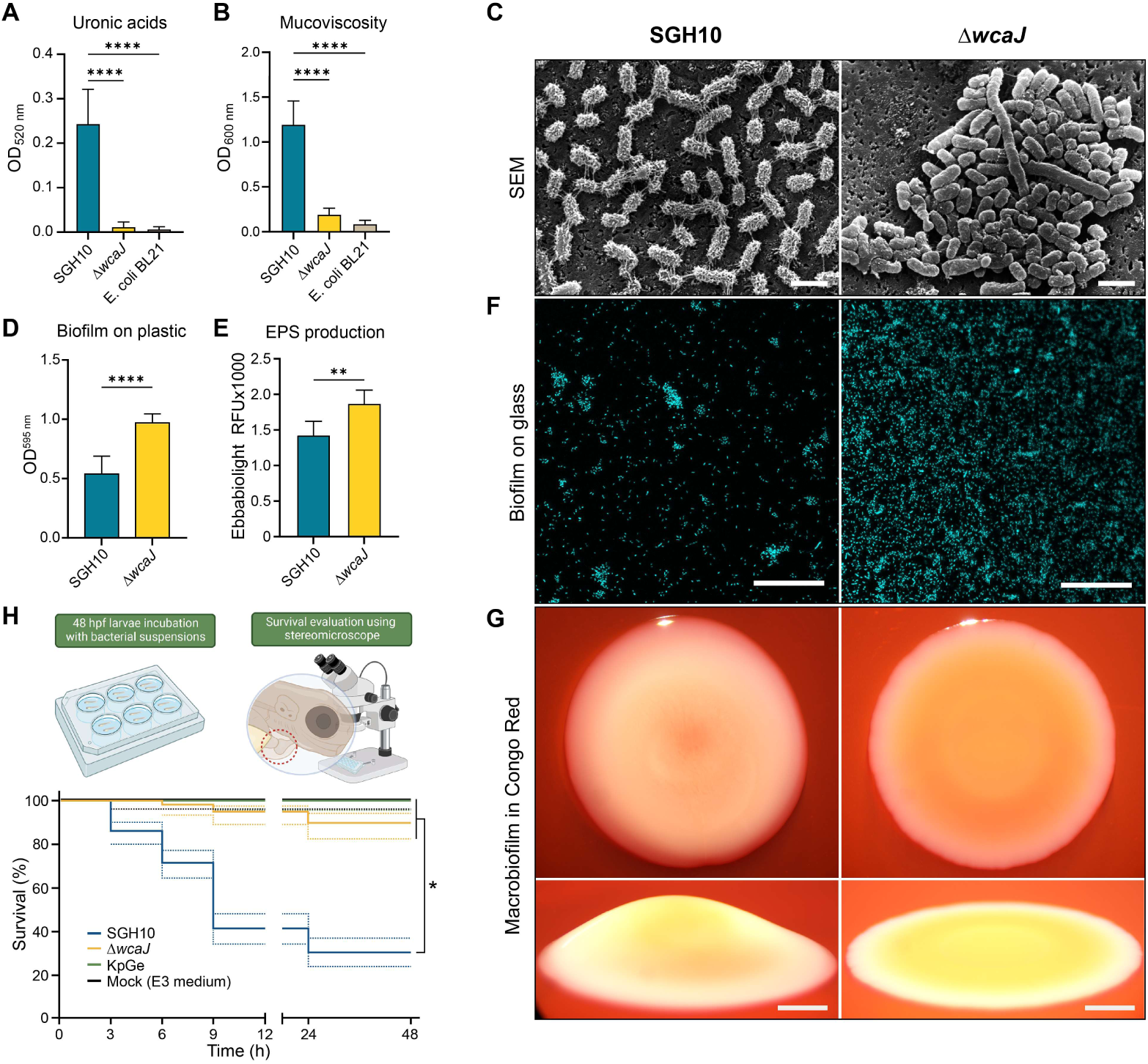
Capsule production, mucoviscosity, biofilm formation, exopolysaccharide (EPS) synthesis, and virulence over zebrafish are affected in the hvKp. Δ*wcaJ* mutant. (A) Capsule production was determined by colorimetric uronic acid quantification. (B) Mucoviscosity was determined by low-speed centrifugation in LB medium, followed by measuring the supernatant’s optical density at 600 nm. (C) Scanning electron microscopy (SEM) images of cells obtained from log-phase bacterial cultures. Scalebar: 2 µm. (D) Biofilm formation on plastic surface, as measured by 0.1% crystal violet (CV) staining after 24 h incubation in a 96-well plate. (E) EPS production measured by EbbaBiolight 630™ fluorescence in colonies grown on agar plates under biofilm-forming conditions. For panels A, B, D, and E, the mean ± standard deviation was plotted (n=5). For statistical analyses, one-way ANOVA was used for CV quantification and Student’s t-test for EPS quantification. ****: P <0.0001. (F) Biofilm formation on a glass surface. DAPI was used for biomass staining. Scalebar: 100 µm. (G) Macrobiofilm on agar media with 40 µg/mL Congo Red. Pictures were taken of the front (upper panels) and lateral (lower panels) views. Scalebar: 2 mm. (H) Kaplan-Meier survival plot showing the lethality over three days-post-fecundation zebrafish larvae upon injection of different bacterial strains (n=15 larvae per condition). Part of this Figure was created with Biorender.

Concomitant with capsule loss, the Δ*wcaJ* mutant exhibited increased biofilm formation over an abiotic (plastic) surface (Figure 1D), as well as EPS production (Figure 1E), suggesting a compensatory shift toward enhanced surface attachment. This phenotype was consistent with confocal microscopy images, which depict a substantial rise in the *ΔwcaJ* bacterial population adhering to glass, compared to the wt strain (Figure 1F). Also, the macrocolony biofilm structure was altered in the mutant strain (Figure 1G). Altogether, these data support that the Δ*wcaJ* mutant has critical defects in structuring the capsule and extracellular matrix, reported to be key traits of hypervirulent strains.

Finally, we aimed to explore the *in vivo* virulence of hvKp and the *wcaJ* mutant in an animal model other than mice. For this purpose, we leveraged the zebrafish larvae model, previously shown to be suited for evaluating the virulence of cvKp (Marcoleta et al., 2018). We assessed the lethality of hvKP and the Δ*wcaJ* mutant over zebrafish larvae by performing static immersion in bacterial suspensions (5×10^8^ CFU/mL). We used the *K. pneumoniae* KpGe strain as a negative control previously shown to be avirulent over zebrafish larvae (Marcoleta et al., 2018). After 24h, hvKp caused roughly 70% lethality, showing significant differences with zebrafish larvae exposed to Δ*wcaJ* and KpGe strains (10% and 0% lethality, respectively) (Figure 1H). These results align with the avirulent phenotype of this mutant in mice (Tan et al., 2020).

Collectively, these results validate the attenuated phenotype of the *wcaJ* mutant previously reported, extending it to a non-mammal vertebrate host. Although zebrafish larvae are a suitable surrogate host model for *K. pneumoniae*, they still require specialized and costly facilities for husbandry and maintenance. In this line, we have previously demonstrated that *D. discoideum* is a simpler and robust host for studying *K. pneumoniae* virulence. Therefore, we aimed first to validate the use of this model in hvKp.

### *D. discoideum* is a suitable model to study hypervirulent *K. pneumoniae*

We initially used two classical assays described previously, relying on simple macroscopic observations: the social development and the predation resistance assays. The first one assesses the progression of the amoeba multicellular stages until culmination with the formation of the fruiting bodies. Previous studies have demonstrated that virulent bacteria, including *Salmonella* Typhimurium, *P. aeruginosa*, and *K. pneumoniae*, can delay or impede the *D. discoideum* social development (Bravo-Toncio et al., 2016; Dunn et al., 2018; Rojas et al., 2024; Varas et al., 2018). In contrast, attenuated or avirulent bacteria permit the rapid progression of the social cycle, which involves at least three stages: aggregation, including the formation of a phagocytosis plaque and subsequent cluster formation; elevation, involving the formation of slugs, fingers, and Mexican hats; and culmination, comprising the formation and maturation of fruiting bodies (Figure 2A).

**Figure 2.**
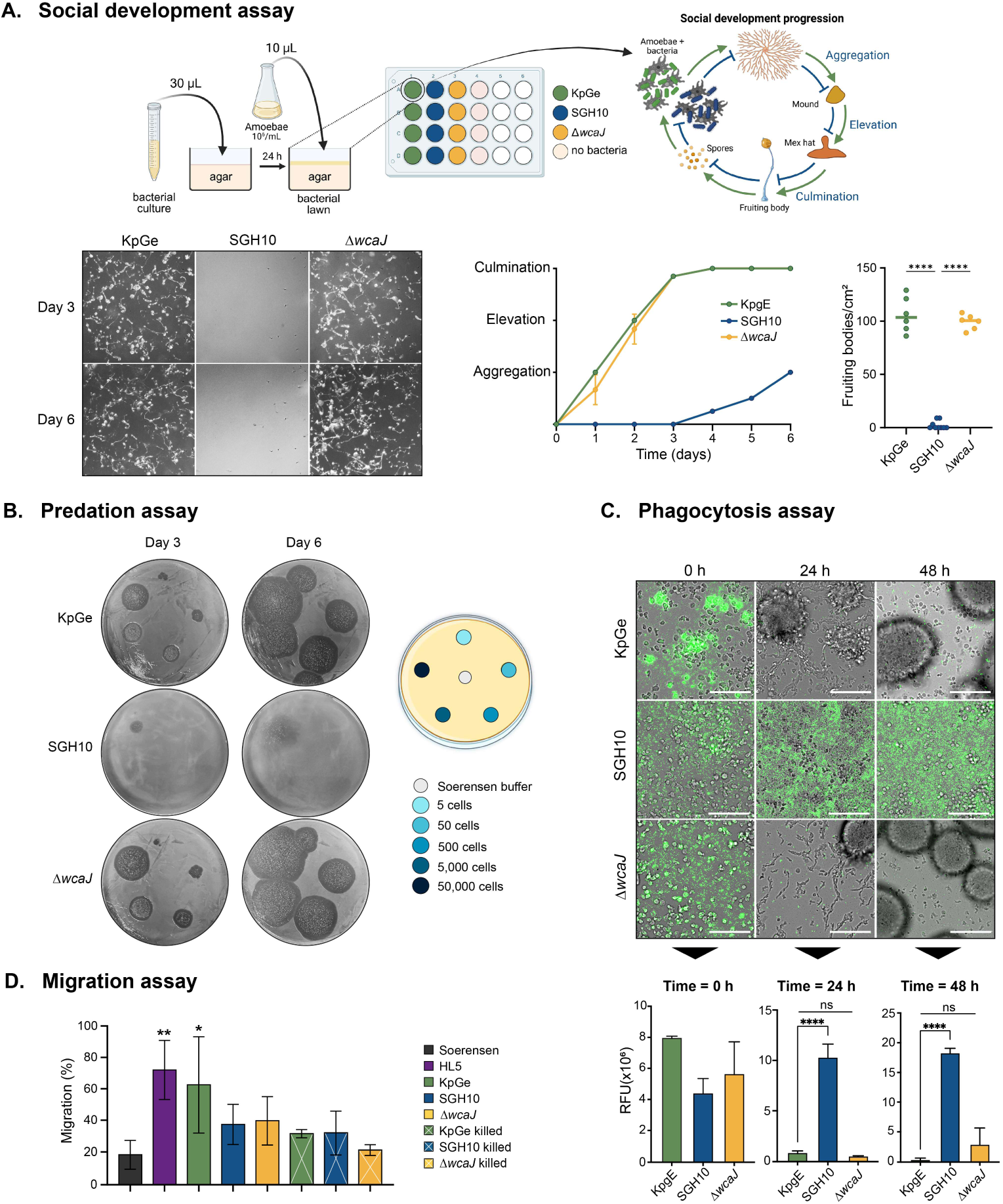
Validation of virulence assays for hypervirulent *K. pneumoniae* based on *D. discoideum* phagocytosis and social development. (A) For social development assays, 30 µL of overnight bacterial cultures were inoculated in a 24-well plate on N agar and incubated for 24 h at 23 °C to generate a lawn. *D. discoideum* suspensions were adjusted to 10^6^ cells/mL. A 10 µL drop of the suspension was inoculated on each lawn. Representative images of days 3 and 6 are presented. The social development of the amoeba was followed for six days, counting the number of fruiting bodies on the third day (D). The avirulent strain KpGe was used as a control permissive for development. (B) For predation resistance assays, overnight bacterial cultures were seeded onto an HL5 agar plate and incubated for 24 h at 23°C to generate a bacterial lawn. *D. discoideum* was cultured in HL5 medium, and serial dilutions were prepared in Sorensen buffer. Subsequently, 5 μL of each *D. discoideum* suspension were applied onto the bacterial agar plates and incubated at 21°C for 6 days. Phagocytosis plaque formation was examined on days 3 and 6. (C) 48 h phagocytosis assay followed by time-lapse microscopy of bacterial strains expressing GFP. Scale, 100 μm. Fluorescence quantification was performed for three biological replicates. One-way ANOVA tests were performed for statistical analyses. **** p < 0.00001. Part of this Figure was created with Biorender. (D) Percentage migration in Trans well assays. The differences were assessed against the basal migration condition (Soerensen buffer). * p < 0.05, ** p < 0.01.

The avirulent strain KpGe, generally used as food to support *D. discoideum* growth, permitted normal social development, reaching culmination between days 2 and 3, and forming, on average, 110 fruiting bodies/cm^2^ on the third day. In contrast, hvKp showed a significant delay, with little or no formation of multicellular structures within the 6-day study period. Only the aggregation phase was observed halfway through day 6, with an average of 2.3 fruiting bodies/cm^2^ observed on the third day. Conversely, the Δ*wcaJ* mutant allowed a normal social development culminating on the third day, with an average of 99 fruiting bodies/cm^2^, resembling the avirulent strain. Representative stereomicroscope photographs are shown in Figure 2A.

The predation assay focuses on *D. discoideum* feeding on bacteria. When even a small number of amoeba cells (i.e., five cells) are inoculated into a lawn of avirulent bacteria, visible phagocytosis plaques form by the second or third day of cultivation. In contrast, when amoebae are cultured on a lawn of virulent bacteria, no visible phagocytosis plaques are formed unless the *D. discoideum* inoculum is increased to some extent. Previous studies have shown that if phagocytosis plaques are formed with 500 or fewer *D. discoideum* cells, the bacteria exhibit susceptibility to predation, indicating reduced or no virulence (Filion and Charette, 2014; Paquet and Charette, 2016).

We recorded the results on the third and sixth day of the assay. The avirulent strain KpGe was used as an edible control, where phagocytosis plaques formed at the lowest concentrations evaluated (5 amoeba cells) (Figure 2B). hvKp strongly resisted amoebae predation and impeded the formation of phagocytosis plaques even with a 50,000-cell inoculum, consistent with a high level of virulence. In contrast, the Δ*wcaJ* mutant was readily phagocytosed and permitted full social development of *D. discoideum*, exhibited a similar behavior to KpGe regarding the size of the plaques in each inoculum for each of the recorded days.

To further support our findings at the microscopic level, we conducted a 48-h time-lapse microscopy experiment to observe the phagocytosis by *D. discoideum* of bacteria constitutively expressing the green fluorescent protein GFP from the pBBR1-GFP plasmid (Porte and Teyssier, 2018). Amoebae were infected with an MOI of 10, and phagocytosis was monitored at 0, 24, and 48 h post-infection (hpi) (Figure 2C). For KpGe, we observed streams of amoebas moving towards aggregation centers within the first 24 h, followed by abundant multicellular aggregation bodies by 48 hpi. The number of bacteria present significantly decreased within the first 24 h, as evidenced by the reduced fluorescence intensity in the green field. In the case of hvKp, no cell aggregation was observed. Moreover, by 48 h, we observed a notorious bacterial proliferation. In contrast, the Δ*wcaJ* mutant showed aggregation centers within the first 24 h and numerous multicellular clusters after 48 h, with a concomitant reduction in bacterial cells.

We used Transwell devices to study possible differences in migratory behavior in response to hvKp and attenuated strains, either alive or heat-inactivated (Figure 2D). The Soerensen buffer condition was considered as a basal migration without an attractant, while the HL5 nutritive medium was used as a positive migration control. Amoebae recruitment required viable bacteria, as non-viable preparations triggered minimal movement. Migration was highest toward the nutritive medium and the edible strain KpGe, whereas hvKp and Δ*wcaJ* showed a migration comparable to the control. Hence, the absence of a capsule did not enhance attraction, arguing against the capsule per se as a chemotactic cue for *Dictyostelium*. Consistent with this, the migrated fraction showed higher viability than the non-migrated fraction, supporting the idea that viability-linked signals, rather than capsular features, drive amoebae engagement.

These results validate the use of this host, particularly features related to phagocytosis, social development, and cell motility to study hvKp virulence. Moreover, they support the central role of the capsule, also in hvKp, as a virulence factor for the evasion of phagocytic cells, including *D. discoideum*.

### Tuning the *D. discoideum* model to distinguish between avirulent, virulent, and hypervirulent *K. pneumoniae* strains

Building on this model, we next aimed to use the amoeba to discriminate among avirulent, virulent, and hypervirulent *K. pneumoniae*. For this, we systematically analyzed the amoeba-bacteria interaction using refined and quantifiable versions of the methodologies previously described. These enhancements allowed us to detect nuanced differences in host-pathogen interaction outcomes that reflect distinct bacterial virulence profiles.

First, we conducted time-course assays to monitor predation over bacterial lawns and the social development dynamics across six days, but now incorporating quantitative measurements. Moreover, we also assayed mixtures of cvKp or hvKp with edible (avirulent) *K. pneumoniae*, addressing whether bacterial virulence affects the feeding capacity of the amoeba, providing additional readouts of each assay.

In the predation assays, we quantified during six days the speed of the feeding front of the phagocytosis plaques formed upon inoculation of different amounts of amoeba cells, namely, 25-25000 cells. The feeding front corresponds to the spatially organized zone at the leading edge of migrating amoeba cells, where active phagocytosis of bacteria occurs, characterized by coordinated chemotaxis, cell polarization, and localized bacterial clearance (Bozzaro and Eichinger, 2011). In this case, we used lower amoeba inocula compared to the classical predation assay to observe a more gradual development. As expected, *D. discoideum* readily predated the KpGe and the Δ*wcaJ* strains, forming a well-defined and homogeneous feeding front and numerous fruiting bodies at day six, when 25000 amoeba cells were inoculated (Figure 3A). In contrast, hvKp remained resistant to amoebal predation, with only a faint plaque formation observed even after extended incubation. In the case of cvKp, we observed intermediate behavior, with plaque formation and expansion evident over time, but showing a more diffuse and disorganized feeding front, with the formation of fewer fruiting bodies compared to the edible strain.

**Figure 3.**
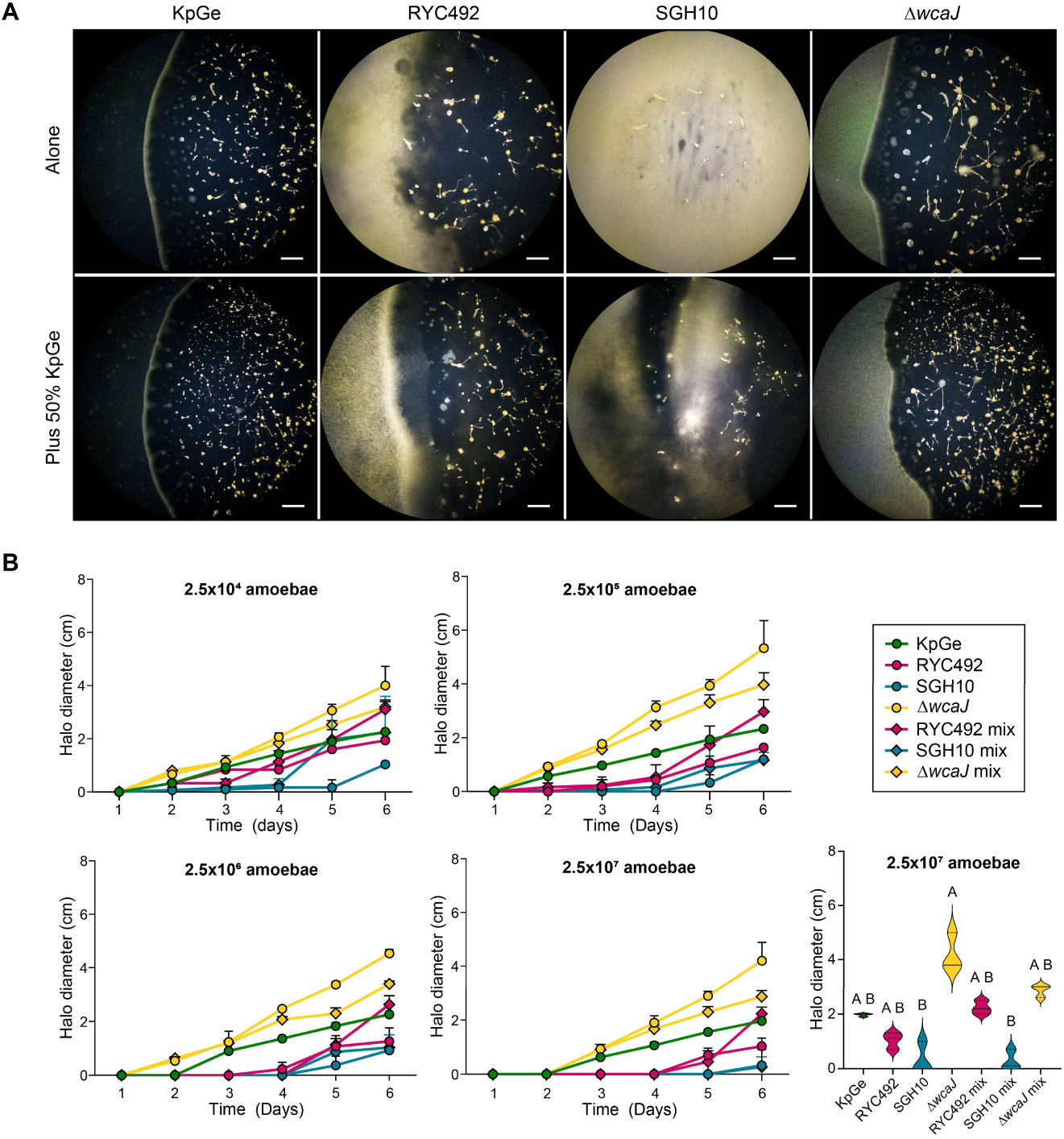
Quantification of *D. discoideum* predation over avirulent, virulent, and hypervirulent *K. pneumoniae.* (A) Representative photographs of the feeding front six days post amoeba inoculation. Scalebar: 400 µm. (B) Phagocytosis plaque diameter over time.

The addition of 50% of edible bacteria favored amoeba predation of the virulent RYC492 strain, leading to the formation of more fruiting bodies and a less diffuse and more organized feeding front. Furthermore, phagocytosis plaques were observed for the SGH10 mixture, although with a severely disrupted feeding front organization (Figure 3A).

The plaque observations over time were used to quantify the feeding front advance speed (Figure S2) and the phagocytosis plaque diameter (Figure 3B) observed for different bacterial strains and amoebae inocula. Both measurements showed a clear separation between virulent and hypervirulent phenotypes.

Social development assays further confirmed this stratification. In this more quantitative version, we counted the different multicellular structures present in a determined area on each day. When grown on a lawn of the edible KpGe strain, roughly 100 mature fruiting bodies per mm^2^ were observed after six days (Figure 4). A similar behavior was observed with the attenuated mutant Δ*wcaJ*. In contrast, amoebae exposed to hvKp and cvKp remained mostly undifferentiated, stalling in the aggregation phase, forming few or no multicellular structures and failing to culminate. Remarkably, the cvKp mixture was more permissive for amoeba development, evidenced by the presence of more early-stage multicellular structures and immature fruiting bodies on day six.

**Figure 4.**
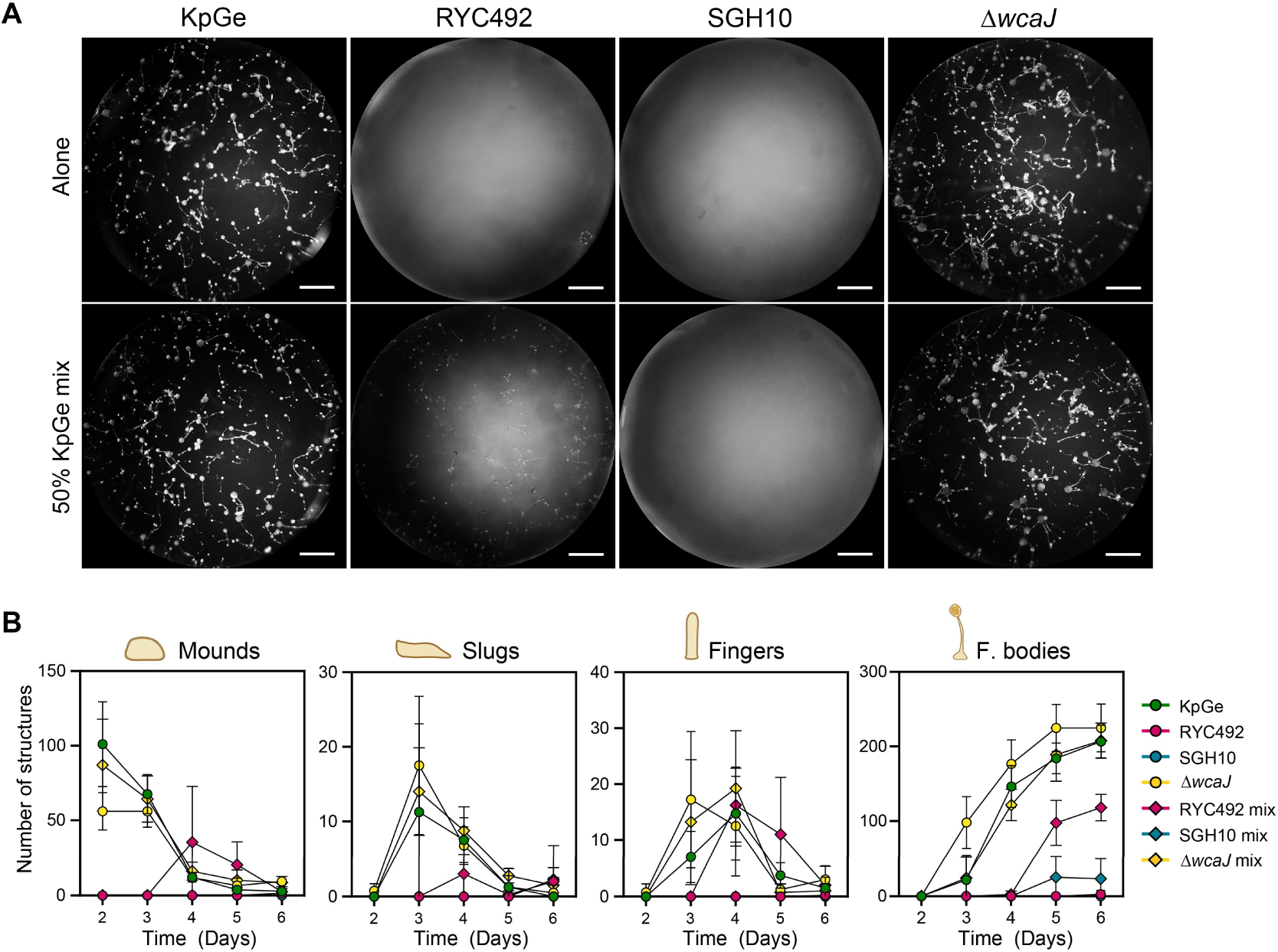
Quantification of multicellular structures during *D. discoideum* social development prove differences between virulent and hypervirulent *K. pneumoniae*. (A) Density of multicellular structures over time. Four independent experiments were made. The bar plots show the average and standard deviation for each condition. (B) Representative pictures of *D. discoideum* social development on the sixth day after exposure to different bacterial strains. Scalebar: 500 µm.

These findings demonstrate that *D. discoideum* can distinguish edible, virulent, and hypervirulent *K. pneumoniae* strains and that this discrimination can be quantified across a battery of readouts, reinforcing its utility as a genetically tractable model to dissect virulence mechanisms in this pathogen. Notably, restoring susceptibility and social development upon inactivating capsule biosynthesis (Δ*wcaJ*) supports the notion that capsule-associated traits drive hypervirulence and resistance to phagocytic clearance in amoebae.

### Single-cell tracking reveals changes in amoeba motility upon challenge with virulent, hypervirulent, or edible *K. pneumoniae*

As shown previously, different bacterial strains induce different *D. discoideum* migratory responses. Thus, we aimed to develop an improved cell migration assay based on time-lapse live cell tracking, using automated fluorescence microscopy to monitor the motility and morphology of *D. discoideum* cells expressing GFP while interacting with bacteria of different virulence degrees (Figure 5A). This way, we explored *K. pneumoniae* virulence effects on amoeba motility and cell integrity as additional readouts for evaluating and finely distinguishing *K. pneumoniae* virulence based on this host model. First, we focused on two parameters: 1) persistence (how long individual spots/cells remain visible across frames); and 2) displacement (the straight-line distance between the first and last spot in a track).

**Figure 5.**
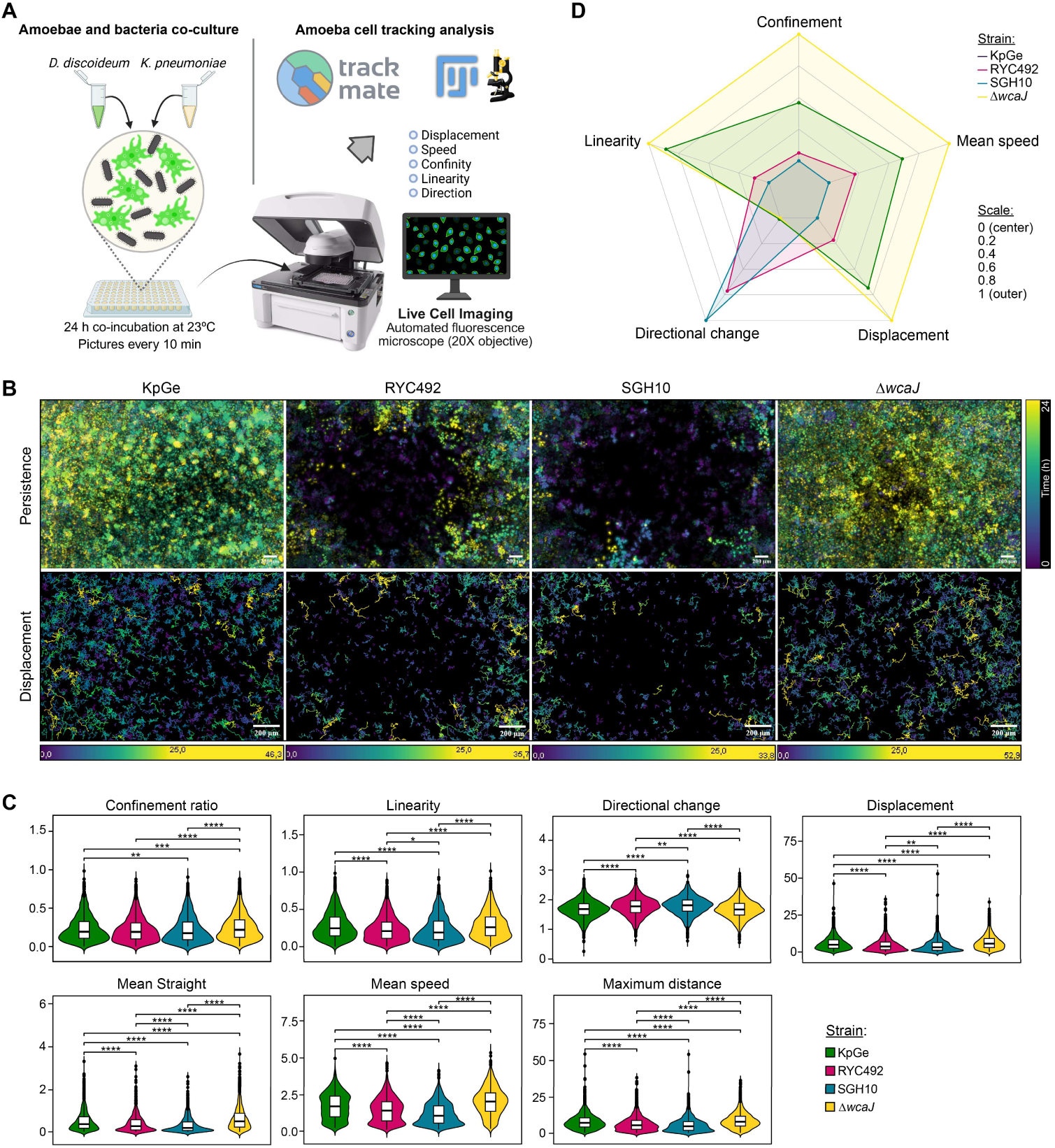
Time-lapse and single-cell tracking assays during bacteria-amoeba interaction revealed impaired amoebae motility linked with *K. pneumoniae* virulence. (A) Schematic representation of time-lapse fluorescence microscopy assays. Partly created with Biorender. (B) Overlay of all the *D. discoideum* cells tracked during 24 h, colored by persistence (upper row) or displacement (lower row). (C) Quantification of the cell-tracking parameters. All variables were tested for normality and homogeneity of variances before group comparisons. As none of the datasets followed a normal distribution, non-parametric analyses were performed using the Kruskal–Wallis test, followed by Dunn’s post hoc test for multiple comparisons. All analyzed parameters, including confinement ratio, linearity, directional change, displacement, mean straightness, mean speed, maximum distance, and total distance, showed significant differences among groups (p < 0.05). Significance thresholds were defined as follows: p < 0.05 (*), p < 0.01 (**), p < 0.001 (***), p < 0.0001 (****). Error bars indicate the median and interquartile range.

Visual overlays of tracked amoebae over 24 hours revealed clear behavioral distinctions after exposure to bacterial strains with varying virulence (Figure 5B-5C, Movies S1-S4). Amoeba cells exposed to the edible strain and Δ*wcaJ* displayed long-lasting permanence and active migration across the field. In contrast, cells incubated with hvKp exhibited highly restricted movement, in a sort of paralysis phenotype likely induced by hvKp. Meanwhile, the exposure to cvKp caused intermediate effects.

Observation and quantification of the GFP intensity over time revealed that, when exposed to KpGe, amoeba cells rapidly phagocytosed bacteria, having populated all the well after 18 h. In contrast, amoebae progressively lost their fluorescence after challenge with hvKp and, to a lesser extent, cvKp, suggesting disruption of some amoeba cells (Figure 6).

**Figure 6.**
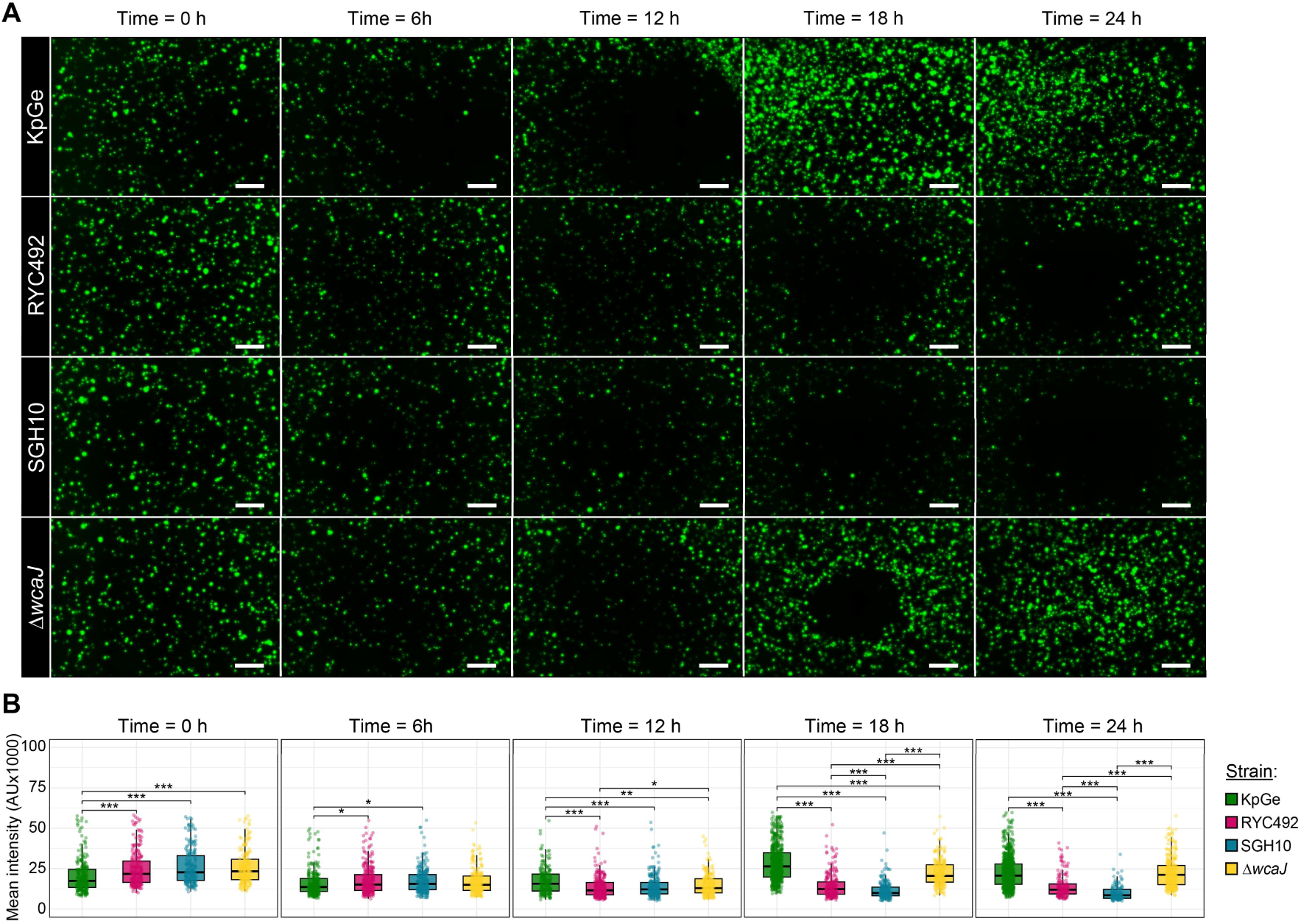
Different degrees of *K. pneumoniae* virulence correlate with amoeba cell disruption. (A)Time-lapse fluorescence imaging of *D. discoideum* cells expressing GFP, after being mixed with *K. pneumoniae* strains. Scalebar: 50 µm. (B) Quantification of mean fluorescence intensity over time. Mean fluorescence intensity values were tested for normality and homogeneity of variances prior to group comparisons. Since the datasets did not follow a normal distribution, non-parametric analyses were performed using the Kruskal–Wallis test, followed by Dunn’s post hoc test for multiple comparisons. Significance thresholds were defined as follows: p < 0.05 (*); p < 0.01 (**); p < 0.001 (***); p < 0.0001 (****). Error bars indicate the median and interquartile range.

In the time-lapse fluorescence images (Figure 6A, Movies S5-S12), *D. discoideum* cells exposed to the different *K. pneumoniae* strains exhibited distinct spatial dynamics over time. Notably, in wells containing hypervirulent SGH10 or its capsule mutant Δ*wcaJ*, amoebae progressively cleared from the central field, accumulating instead at the periphery after 12–18 h. This apparent “exclusion zone” suggests that cells at the center may experience physical or mechanical constraints that limit their movement or survival. Such behavior evokes the active droplet-like patterning described by Chubb et al. (Ford et al., 2025), where collective motility and local depletion create ring-like arrangements within cell populations. In this context, the observed peripheral migration could reflect self-organization along chemotactic or mechanical gradients, with central regions becoming depleted due to cell death, adhesion changes, or mechanical interference, while peripheral cells remain motile and responsive.

These results suggest that hvKp triggers rapid amoebae paralysis, whereas attenuated or edible strains allow amoebae to retain motility and dynamic shape changes. This single-cell resolution approach complements traditional developmental and phagocytic assays, reinforcing the value of *D. discoideum* as a morpho-functional model for early-stage host-pathogen interaction studies.

### Integrating multivariate data for virulence differentiation

To integrate and compare the performance of multiple phenotypic assays, principal components analysis (PCA) was used as an exploratory proof-of-concept visualization to summarize correlated, large-effect phenotypes measured across predation, social development, and cell movement, including quantitative variables from all these traits. The first two principal components accounted for over 90% of the total variance, effectively capturing the distribution of strain-specific phenotypes (Figure 7).

**Figure 7.**
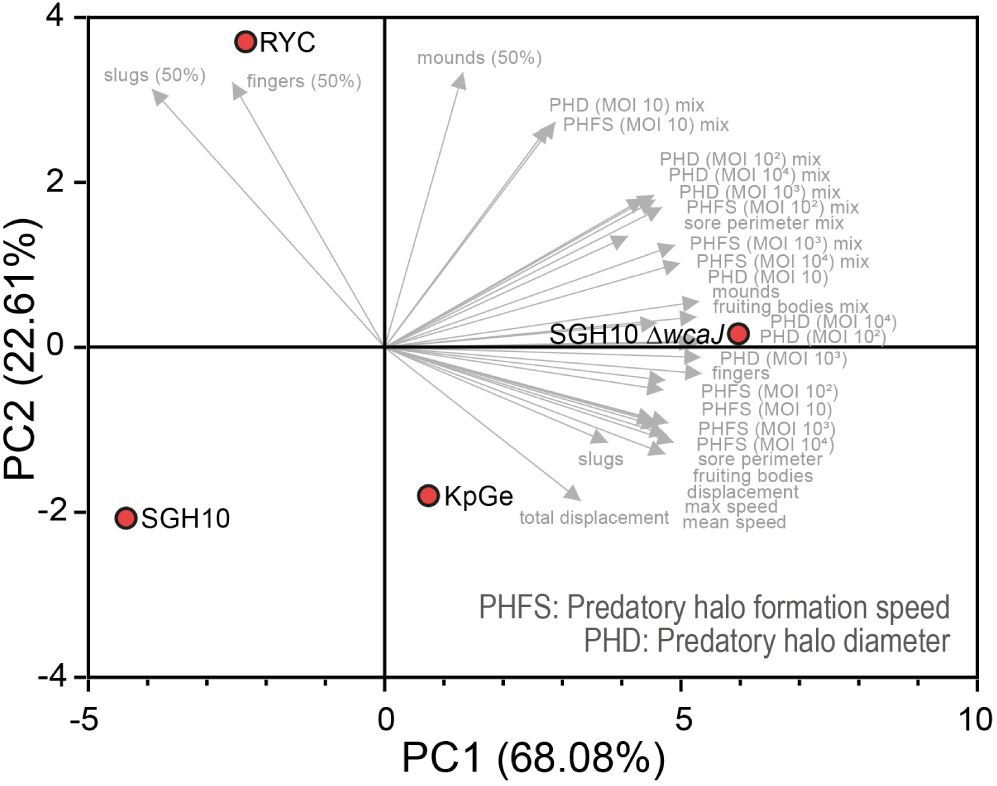
Principal component analysis (PCA) based on multiple measurements during the interaction of *D. discoideum* and *K. pneumoniae* strains of varying virulence degrees. The conditions indicating “mix” are mixtures of the respective strain with 50% KpGe.

The first component (68%) clearly separated virulent versus edible strains, supported by most of the variables included. The second one differentiated cvKp from hvKp, where the latter correlated negatively with variables such as the number of fruiting bodies and other multicellular structures, predatory halo diameter and formation speed, displacement, and velocity, among others. cvKp clustered apart from hvKp mainly by permitting amoeba development when mixed with the edible strain KpGe, reflecting the hvKp higher relative virulence. Strikingly, the capsule mutant Δ*wcaJ*, although differing by only one gene from the hvKp strain, showed the strongest association with most of the variables, seeming to be even more edible than the KpGe strain, confirming its highly attenuated phenotype.

Together, these multivariate analyses support the use of *D. discoideum* as a robust phenotypic biosensor capable of distinguishing edible, virulent, and hypervirulent *K. pneumoniae* strains through quantitative behavioral and morphological signatures. This approach enhances the resolution of traditional host-pathogen interaction assays and can be adapted for high-throughput screening. Forthcoming analyses with cross-validated metrics and a larger set of strains are required to validate this multivariate integration and define quantitative thresholds and performance metrics.

## DISCUSSION

The ability to functionally characterize the virulence of *K. pneumoniae* strains in a rapid, scalable, and mechanistically informative manner remains critical in clinical microbiology and antimicrobial resistance surveillance, especially for hypervirulent and convergent strains. This study demonstrates that the social amoeba *D. discoideum* provides a powerful platform to phenotypically differentiate edible, virulent, and hypervirulent *K. pneumoniae* strains. Using a combination of predation, social development, and real-time single-cell tracking assays, we show that *D. discoideum* recapitulates key features of the host-pathogen interaction in a non-mammalian system, particularly those associated with the capsular polysaccharide, immune evasion, and phagocytosis resistance.

Recent findings reinforce the central role of the capsule in hypervirulence and its position as one of the most critical virulence factors in *K. pneumoniae*, which shows phase variation to balance immunogenicity, stress resistance, adhesion, and horizontal gene transfer (Liu et al., 2025; Paczosa and Mecsas, 2016; Wei et al., 2025). Furthermore, CPS overproduction is a hallmark of hypervirulent strains, often driven by the *rmpADC* operon, which enhances mucoviscosity and promotes metastatic dissemination (Ovchinnikova et al., 2023; Walker et al., 2019; Zhu et al., 2021). Moreover, recent transcriptomic and RIL-seq (RNA Interaction by Ligation and Sequencing) analyses have revealed that capsule regulation in hvKp is tightly controlled by small RNAs and global regulators, further underscoring its centrality in virulence expression (Goh et al., 2024). Our study shows that deletion of a single gene, Δ*wcaJ*, encoding the initiating glycosyltransferase for capsule biosynthesis, is sufficient to abolish hypervirulent traits in SGH10, yielding a phenotype indistinguishable from edible strains. This aligns with prior murine model observations (Tan et al., 2020) and extends them to amoebal and zebrafish hosts. Collectively, these findings support the capsule as a phenotypic anchor for hypervirulence and a mechanistic driver of host evasion across diverse infection models.

Interestingly, despite capsule loss, Δ*wcaJ* exhibited increased biofilm formation and EPS production, consistent with prior reports in other *K. pneumoniae* strains (Singh et al., 2019). This shift toward surface attachment may enhance horizontal gene transfer, as mutations in *wcaJ* have been linked to increased recombination frequency of blaKPC-2 plasmids (Wang et al., 2022). Such changes could facilitate the exchange of resistance and virulence determinants, contributing to the evolution of convergent strains. Capsule disruption has also been associated with increased epithelial invasion and persistence in urinary tract infections (Ernst et al., 2020), while mutations in *wzc*, involved in capsule polymerization and translocation, can lead to hypercapsulation and metastatic dissemination (Teng et al., 2025). These findings underscore the capsule’s dual role in immune evasion and ecological adaptation, and highlight the need for functional assays that can resolve these phenotypic nuances.

The present study provides a quantitative framework for virulence detection using *D. discoideum* as a living biosensor, bridging the gap between genomic prediction and functional pathogenicity. By integrating automated imaging, behavioral metrics, and phenotypic clustering, this approach enables rapid discrimination of hypervirulent *K. pneumoniae* strains while maintaining scalability and reproducibility. Incorporating such phenotypic biosensing systems alongside genomic and antimicrobial-resistance profiling will be critical for diagnosing convergent pathotypes, which represent a growing clinical and epidemiological threat.

A significant advantage of this model lies in its capacity to discriminate virulence levels based on amoebal behavior and developmental progression. In agreement with prior reports (Marcoleta et al., 2018; Rojas et al., 2024), cvKp and especially hvKp resisted phagocytic clearance and blocked multicellular development in *D. discoideum*. We extended these observations by incorporating dynamic quantification of phagocytosis plaque expansion, fruiting body formation kinetics, and amoebae motility. The integration of these data confirmed that capsule biosynthesis, abolished in the Δ*wcaJ* mutant, is a key determinant of virulence and is sufficient to shift the phenotype from hypervirulent to one resembling edible commensals. Importantly, we introduce live cell tracking as a novel phenotypic assay to quantify amoebal paralysis, a functional outcome of virulence that correlates with evasion of phagocytosis. Amoebae exposed to hvKp rapidly adopted a rounded, immobilized morphology with high circularity and reduced displacement, consistent with functional paralysis. In contrast, strains such as Δ*wcaJ* or KpGe supported active amoebal motility. These behavioral signatures illustrate how *D. discoideum* can capture complex virulence traits at both the population and cellular levels.

From a translational perspective, the scalability and genetic tractability of *D. discoideum* offer significant advantages over traditional vertebrate models. This model supports large-scale screening of convergent evolution between resistance and virulence traits by enabling the parallel phenotypic assessment of a large number of clinical isolates. Moreover, the ability to map phagocytosis resistance, capsule function, and developmental interference in a single, low-cost platform opens the door to studying hypervirulence evolution without reliance on murine infection models.

While our study focused on a limited number of *K. pneumoniae* isolates, this was a first step to establish and calibrate the assay framework using well-characterized strains. A broader strain diversity is required to capture the clinical and ecological spectrum of virulence phenotypes fully. Further studies expanding the panel to include a wider array of isolates spanning different sequence types, capsule loci, and resistance profiles are warranted. This will allow us to define quantitative thresholds for virulence classification and begin assessing the diagnostic performance of the platform.

While our assays robustly captured capsule-dependent virulence traits, particularly through the SGH10/Δ*wcaJ* comparison, it remains to be determined whether the *D. discoideum* platform can resolve other hypervirulence determinants beyond the capsule. Notably, other virulence plasmid-encoded factors, including siderophore systems such aerobactin (*iuc*) and salmochelin (*iro*), contribute differentially to pathogenesis depending on the infection niche (Lim et al., 2025). Testing mutants in these *loci* across our functional assays would provide valuable insight into the model’s sensitivity and discriminatory power. This is especially relevant for detecting convergent strains, which may harbor combinations of *rmpA*, one or more siderophores, and variable capsule expression. Expanding the assay suite to include such mutants will help define the resolution limits of the amoeba biosensor and its potential for early-stage screening of complex virulence architectures.

Beyond *K. pneumoniae*, the virulence biosensor paradigm demonstrated here can be readily extended to other clinically relevant bacteria, including the ESKAPE group. Recent studies have revealed hypervirulent phenotypes emerging in non-classical species, such as *P. aeruginosa* strains co-expressing the T3SS effectors ExoS and ExoU (Song et al., 2023), or hypervirulent lineages of *E. coli* associated with invasive extraintestinal infections (Ba et al., 2024). The adaptability, genetic tractability, and quantitative sensitivity of *D. discoideum* make it a versatile surrogate for dissecting virulence mechanisms across diverse pathogens, opening the way to biosensor-driven platforms for comparative virulence assessment, antivirulence screening, and real-time environmental surveillance. Furthermore, the suitability of *D. discoideum* as a host for virulence studies was validated in several species besides *K. pneumoniae*, including *P. aeruginosa*, *S.* Typhimurium, *E. coli*, and *Legionella pneumophila*, among others.

In conclusion, this work establishes *D. discoideum* as a living analytical platform that bridges microbial pathogenesis and quantitative biosensing. By translating cellular behavioral responses into measurable virulence signatures, *D. discoideum* enables the detection of functional pathogenic traits that are invisible to genomic prediction alone. This integrative approach, linking genotype, phenotype, and antimicrobial susceptibility, provides a scalable and ethical framework for the discovery, monitoring, and comparative evaluation of virulent and multidrug-resistant pathogens. As emerging technologies continue to merge cell biology, automated imaging, and computational analytics, *D. discoideum*-based biosensors hold great promise for advancing precision infection biology and real-time surveillance of microbial threats across clinical and environmental settings.

## METHODS

### Bacterial strains and plasmids

*E. coli* and *K. pneumoniae* strains were routinely grown at 37°C in LB broth (10 g/L Tryptone, 5 g/L Yeast Extract, 5 g/L NaCl), with 1.5% (w/v) agar added for solid medium. Antibiotic selection was performed using 100 μg/mL Carbenicillin, 50 μg/mL Kanamycin, or 10 μg/mL Tetracycline. The bacterial strains included *K. pneumoniae* KpGe (Lima et al., 2018), used to support the growth of *D. discoideum*; *E. coli* S17 λpir, a laboratory cloning strain used as a donor for conjugation; *K. pneumoniae* SGH10, isolated from a human liver abscess in 2014 and characterized as hypermucoviscous and hypervirulent (Lam et al., 2018); and the SGH10 Δ*wcaJ* mutant, a capsule-null mutant lacking the WcaJ glycosyltransferase involved in the first step of capsule polysaccharide synthesis, used as an attenuated strain. Plasmids used included pBBR1-GFP, constitutively expressing GFP with kanamycin resistance, and pR6Ktet-sacB, a vector for gene deletion containing a tetracycline resistance cassette and *sacB* for promoting the second recombination event.

### *K. pneumoniae* site-directed mutagenesis

We followed the pipeline below to acquire mutant strains with a deleted gene of interest. First, we amplified ∼1000 bp fragments upstream and downstream of the target gene from genomic DNA templates using Q5 high-fidelity DNA polymerase (New England Biolabs). We assembled them into the vector pR6Ktet-SacB using NEBuilder® HiFi DNA Assembly Master Mix (New England Biolabs). The plasmids were then introduced into *K. pneumoniae* SGH10 through conjugation with the *E. coli* donor strain S17-1λpir. Transconjugants were selected in a medium containing tetracycline (10 µg/mL) and ampicillin (100 µg/mL). Suicide plasmid integration was confirmed by PCR. For sacB-based counterselection, ten colonies positive for plasmid integration were inoculated into LB medium without sodium chloride, supplemented with 20% sucrose, and allowed to grow overnight. Then, an aliquot was used to inoculate 1:100 fresh medium, and let grow overnight. This procedure was repeated until tetracycline-sensitive colonies appeared. Successful deletion of the gene of interest in tet-sensitive colonies was confirmed via PCR. The confirmation of the scar (Δ*wcaJ*) was performed by sequencing the fragments and matching them with the complete genome of the SGH10 WT.

### Scanning electron microscopy (SEM)

To examine potential modifications at the cell surface level, including the capsule and overall morphology, we conducted scanning electron microscopy (SEM), as follows. We prepared a lawn on LB agar from strains frozen at −80°C, then incubated for 24 h at 37°C. Using a sterile swab, we collected enough bacteria and resuspended them in PBS 1X. We adjusted the OD_600nm_ to 2.8 (10^9^ CFU/ml) and fixed 500 µL of the suspension with 2.5% glutaraldehyde and 0.15% ruthenium red to improve the capsule visualization contrast. Samples were subjected to critical point drying and gold coating and subsequently observed in a high vacuum Zeiss EVO M10 scanning electron microscope.

### Quantification of Biofilm Biomass and EPS and Visualization Using Confocal Microscopy

Biofilm formation was assessed using a previously described protocol (Chen et al., 2020), with some modifications. Briefly, *K. pneumoniae* strains were grown overnight in LB broth, then diluted 1:100 in fresh LB broth. Next, 100 μL of each dilution was added to a 96-well polystyrene microtiter plate and incubated at 37°C for 24 h. After removing planktonic cells, wells were washed twice with sterile water and stained with 150 μL of 0.1% crystal violet for 30 min. Then, wells were rinsed twice with sterile water, and the stained biofilms were solubilized with 95% ethanol. Biofilm formation was quantified by measuring the OD_595nm_ using a TECAN Infinite 200 pro plate reader.

To quantify EPS components such as curli and cellulose, the Ebbabiolight 680 probe was used, as it binds to these components without influencing biofilm formation (Choong et al., 2021). EbbaBiolight 680 was diluted in a growth medium at a 1:1000 ratio for the biofilm assay. Then, the supplemented growth medium was inoculated with the bacterial culture. Next, the wells of a 96-well plate were filled with 100 μL of the inoculated medium. The unused wells were filled with sterile water to prevent drying during incubation. Fluorescence was measured at 540ex/680em in a TECAN Infinite 200 Pro plate reader. Each sample was measured in triplicate, and we calculated averages of the absorbance values for analysis.

To evaluate biofilm formation on a glass surface using confocal microscopy, an overnight culture was diluted 1:100 in the LB broth medium to observe the biofilm under confocal microscopy. Subsequently, a slide was submerged in 10 mL of the 1:100 culture inside a 50-mL conical tube and incubated at 37°C for 24 h. The slide was washed three times with nanopure water and incubated for 30 minutes with 5 µg/mL DAPI to stain DNA. The biofilm was visualized using an LSM Zeiss 710 confocal microscope, with a 358ex/461em laser.

### Capsular polysaccharides extraction and quantification

Capsular polysaccharides (glucuronic acids) were quantified using a modified version of the protocol described previously (Favre-Bonte et al., 1999). In a 2 mL tube, 500 µL bacterial PBS suspension (OD_600nm_ = 4) with 100 µL of 1% Zwittergent 3-10 in 100 mM citric acid (pH = 2) were mixed and incubated at 50°C for 20 min. After incubation, the mixture was centrifuged at 18000g for 5 min. 300 µL of supernatant were transferred into a new tube, and 1.5 mL of absolute ethanol were added, incubating at 4°C for 30 min. Then, the mixture was centrifuged at 18000 x g for 20 min at 4°C, and the ethanol was discarded with a micropipette, followed by allowing the precipitate to dry at 37°C. Subsequently, the precipitate was resuspended in 200 µL of sterile water. Afterward, 100 µL of the suspension were mixed with 12.5 mM of borax in absolute H_2_SO_4_ and cooled for 10 min on ice. Then, it was incubated at 95°C for 5 min and cooled again for 5 min. Finally, 200 µL of the borax-resuspended sample were mixed with 3 µL of 0.15% phenol in 0.5% NaOH, and the absorbance was measured at 520 nm using a 96-well plate reader.

### *K. pneumoniae* hypermucoviscosity assessment

Mucoviscosity was evaluated by low-speed centrifugation (2000 x g) followed by optical density (OD_600nm_) measurement of the supernatant, as performed previously (Tan et al., 2020). Higher OD values indicate greater retention of cells in suspension due to increased mucoviscosity.

### *D. discoideum* growth conditions

We obtained the *D. discoideum* strain AX4 (DBS0302402) from the Dicty Stock Center (Basu et al., 2013; Kreppel et al., 2004) and cultured it using standard protocols (Fey et al., 2007). Briefly, the axenic cultures of *D. discoideum* vegetative cells were maintained at 23°C in SM medium containing 10 g/L glucose, 10 g/L peptone, 1 g/L yeast extract, 1 g/L MgSO_4_ × 7H_2_O, 1.9 g/L KH_2_PO_4_, 0.6 g/L K_2_HPO_4_, and 20 g/L agar. They were grown on a confluent lawn of *Klebsiella aerogenes* DBS0305928 (now renamed as K. pneumoniae KpGe), also obtained from the Dicty Stock Center. Before the assays, amoebae were grown at 23°C with agitation (180 rpm) in liquid HL5 medium, which contained 14 g/L tryptone, 7 g/L yeast extract, 0.35 g/L Na_2_HPO4, 1.2 g/L KH_2_PO_4_, and 14 g/L glucose at pH 6.3.

### *D. discoideum* social development assays

For classical social development assays, we followed a published protocol (Varas et al., 2018). Briefly, overnight cultures of each bacterial strain (30 μL) were homogeneously included per well of a 24-well plate containing N agar (1 g peptone, 1 g glucose, 20 g agar in 1 L of 17 mM Soerensen phosphate buffer) and grown overnight at 23◦C. A drop of a cellular suspension corresponding to 10^4^ *D. discoideum* cells in HL5 was spotted in the middle of each well, and the plates were further incubated at 23◦C for 6 days. The social development of amoebae was observed for 6 days and classified into “aggregation,” “elevation,” and “culmination” phases. A score of “1” indicated amoebae aggregation, “2” for elevated structures, and “3” for fruiting bodies formed. Transitions between phases were scored with half the value of the nearest next stage. The number of fruiting bodies on a 1 cm2 surface was also quantified for each strain under study and was compared with the commensal KpgE strain. Additionally, images of social development were obtained on days 1, 2, and 3 using a Zeiss Stemi 305 stereomicroscope with a total magnification of 20X.

The improved quantitative version of the social development assay was performed as follows. Thirty microliters of an amoeba–bacterium coculture were inoculated onto each well of a 24-well plate containing 1.5 mL of N agar. The coculture was prepared by mixing 100 µL of a bacterial suspension (∼10⁷ cells/mL) with 100 µL of a *D. discoideum* suspension adjusted to ∼10⁶ cells/mL in Soerensen buffer. Images were taken daily (days 2, 3, 4, 5, and 6) using a camera attached to the eyepiece of a stereomicroscope. The developmental structures (mounds, slugs, fingers, and fruiting bodies) were quantified daily, and the sorus area of the fruiting bodies was analyzed using Fiji (ImageJ).

### Predation resistance assays

The classical assay was performed following a previously established protocol (Filion and Charette, 2014). Briefly, bacterial colonies were taken from a −80°C stock to prepare overnight cultures, and 300 µL were then seeded onto a plate using a Digralsky loop to generate a bacterial lawn. The plate was left to dry and incubated for 24 h at 23°C. Meanwhile, *D. discoideum* was cultured in HL5 medium, and serial dilutions were prepared to obtain the following cell concentrations: 500,000, 50,000, 5,000, 500, 50, and 5 cells per 5 µL. The bacterial lawns were then spotted with 5 µL of the serial *D. discoideum* dilutions. The plates were allowed to dry and were incubated at 21°C for 6 days. Plaque formation was visually examined on days 3 and 6. Isolates that did not enable amoebae growth were considered virulent for the amoeba. Bacterial strains that exhibited social development with 500 *D. discoideum* cells or fewer were considered sensitive to predation (Paquet and Charette, 2016).

The improved quantitative version of the predation assay was performed as follows. In 150-mm plates containing 45 mL of SM/5 agar, bacterial suspensions were inoculated to obtain a homogeneous lawn (5 mL inoculated and 4.8 mL removed). Once the surface was dry, 5 µL of *D. discoideum* suspensions at four different concentrations (5×10⁶, 5×10⁵, 5×10⁴, and 5×10³ cells/mL) were spotted at four equidistant points in a clockwise direction. The formation of a clearing halo was monitored daily by measuring its diameter and taking photographs of the plate.

### Phagocytosis assays

A lawn was prepared on LB agar from strains GFP frozen at −80°C and incubated for 24 h at 37°C. Bacteria were collected using a sterile swab and resuspended in Sorensen 1X buffer. Cell density was adjusted to OD^600^ ^nm^ at 10^7^ cells/mL. Axenic cultures of *D. discoideum* were set to a concentration of 10^6^ cells/mL and 1 mL of culture was deposited in 24-well plates. The plate was centrifuged at 600 x g for 10 min and incubated at 23°C for 24 h. After washing each well three times with Sorensen buffer, 1 mL of bacterial suspension was added to reach a MOI of 10. The plate underwent centrifugation at 600 x g for 30 min. The Lionheart FX automated microscope was used to capture sequential images every 10 min for 24 to 48 h. It covered both bright field (BF) and GFP imaging in 4 specific regions of interest per well, all maintained at a consistent temperature of 23°C. The images were analyzed with Gen5 software version 3.08 (Lionheart FX) and ImageJ software version 1.52 (Fiji version).

### Transwell cell migration assays

Migration of *D. discoideum* was evaluated using 24-well Transwell® plates (8 μm pore polycarbonate membranes, Corning) following a modified Boyden chamber protocol (Boyden, 1962). Briefly, 500 μL of chemoattractant or bacterial suspension was placed in the lower chamber, and 100 μL of amoebae suspension (1 × 10⁶ cells/mL) was added to the upper insert, yielding 1 × 10⁵ cells per condition. Plates were sealed and incubated at 23 °C with gentle agitation (50 rpm) for 6 h. After incubation, membranes were removed, the apical side was gently cleaned to eliminate non-migrated cells and stained with 2% crystal violet. Migrated cells were quantified microscopically and complemented by a viability-based phagocytosis halo assay. Three fractions were analyzed, non-migrated (apical), migrated-attached (membrane), and migrated-suspended (basal), by plating each fraction with *K. pneumoniae* KpGe transformed with the pBBR1-mApple plasmid on SM agar and counting phagocytosis halos after 72–96 h. This combined approach enabled quantitative assessment of *Dictyostelium* migration and viability under different bacterial strains.

### Single cell tracking assays

In transparent, flat-bottom 96-well plates, 100 µL of bacterial suspension (∼10⁷ cells/mL) and 100 µL of GFP-expressing *D. discoideum* (∼10⁶ cells/mL) were added to each well. The plate was centrifuged at 2,000 rpm for 5 min to promote cell settling. Time-lapse imaging was performed using the automated microscope Lionheart FX. Brightfield and green fluorescence (excitation filter: 469 ± 35 nm; emission filter: 525 ± 39 nm) images were acquired every 10 min with a 20X objective at a constant temperature of 23°C. The resulting images were processed using Fiji (Schindelin et al., 2012) with the TrackMate plugin (Tinevez et al., 2023) to perform cell tracking.

### Lethality over zebrafish by static immersion assays

To evaluate lethality o<u>v</u>er Zebrafish, an adaptation of the immersion infection protocol developed previously (Varas et al., 2017) was performed. 48 hpf dechorionated Tab5 larvae were used. Bacterial suspensions of *K. pneumoniae* SGH10 and Δ*wcaJ* were prepared on 50 mL tubes with sterile E3 medium, adjusted at OD_600nm_ = 1,4 (equivalent to ∼ 1×10^9^ CFU/mL, according to viable bacteria recount by serial dilution and plating on LB-agar plates). Afterward, using 6-well plates, 10 to 15 larvae were deposited in a well containing 4 mL of sterile E3 and 4 mL of bacterial suspension. Each condition was assayed in duplicates involving 30 individuals, including the control condition only with sterile E3 medium (lacking bacteria).

The plates were incubated at 28°C for 48 h, keeping the respective 12 h light/ 12 h dark photoperiod. Survival was recorded every three hours, the first nine hours post-exposition (hpe), and then recorded at 24 and 48 hpe, removing dead larvae. To evaluate the survival, tactile stimulation was applied to the larvae’s tail, and visualization of the heartbeat was made using a Zeiss Stemi 305 stereomicroscope when confirmation was needed. Kaplan-Meier survival curves were constructed with the data obtained using GraphPad Prism version 8.0.1 software. Statistical analysis was performed with the same software using 2-way ANOVA tests with Bonferroni’s multiple comparison post-test.

## SUPPLEMENTARY INFORMATION

**Figure S1.** Overview of the site-directed mutagenesis strategy based on the suicidal plasmid pR6K-tet-sacB to generate the *K. pneumoniae* SGH10 Δ*wcaJ* deletion mutant.

**Figure S2.** Predation halo formation speed of different amounts of *D. discoideum* cells after being seeded on a lawn of distinct *K. pneumoniae* strains.

**Movies S1-S4.** Time-lapse tracking of *D. discoideum* cells expressing GFP after exposure to avirulent *K. pneumoniae* (S1), cvKp (S2), hvKp (S3), and the attenuated hvKp mutant Δ*wcaJ* (S4). The images were acquired in the Lionheart FX automated fluorescence microscope, using a 20X objective. The traces show the cell displacement, as determined using the TrackMate package and the Fiji software. All the movies were built from images that were taken every 10 minutes over 24 hours, and include the representative images shown in Figure 5B.

**Movies S5-S12.** Time-lapse fluorescence microscopy imaging of GFP-tagged *D. discoideum* cells interacting with avirulent *K. pneumoniae* (S5-S6), cvKp (S7-S8), hvKp (S9-S10), and attenuated hvKp (S11-S12). The movies S5, S7, S9, and S11 show a merge between the bright field and the green fluorescence channels, while movies S6, S8, S10, and S12 show only the GFP channel. All the movies were built from images that were taken every 10 minutes over 24 hours, and include the representative images shown in Figure 6A.

## ACKNOWLEDGEMENTS

We want to acknowledge Nicole Molina for her continuous support of our experimental work. Also, to the Unidad de Microscopía Avanzada (UMA, Facultad de Ciencias, Universidad de Chile) and the project FONDEQUIP EQM180216 (Lionheart FX equipment), who supported the microscopy studies. Ian Perez acknowledges the Beca Doctorado Nacional 21232343 Doctoral scholarship from Agencia Nacional de Investigación y Desarrollo ANID (Chile).

