## Supplementary Figures and Movies for "Expanding the toolbox for hypervirulent *Klebsiella pneumoniae* using the social amoeba *Dictyostelium discoideum* as a virulence biosensor": Supplementary material v141125.pdf

**Figure S1.** Overview of the site-directed mutagenesis strategy based on the suicidal plasmid pR6K-tet-sacB to generate the *K. pneumoniae* SGH10 D*wcaJ* deletion mutant.

**Figure S2.** Predation halo formation speed of different amounts of *D. discoideum* cells after being seeded on a lawn of distinct *K. pneumoniae* strains.

**Movies S1-S4.** Time-lapse tracking of *D. discoideum* cells expressing GFP after exposure to avirulent *K. pneumoniae* (S1), cvKp (S2), hvKp (S3), and the attenuated hvKp mutant D*wcaJ* (S4). The images were acquired in the Lionheart FX automated fluorescence microscope, using a 20X objective. The traces show the cell displacement, as determined using the TrackMate package and the Fiji software. All the movies were built from images that were taken every 10 minutes over 24 hours, and include the representative images shown in Figure 5B.

**Movies S5-S12.** Time-lapse fluorescence microscopy imaging of GFP-tagged *D. discoideum* cells interacting with avirulent *K. pneumoniae* (S5-S6), cvKp (S7-S8), hvKp (S9-S10), and attenuated hvKp (S11-S12). The movies S5, S7, S9, and S11 show a merge between the bright field and the green fluorescence channels, while movies S6, S8, S10, and S12 show only the GFP channel. All the movies were built from images that were taken every 10 minutes over 24 hours, and include the representative images shown in Figure 6A.

A

B

C

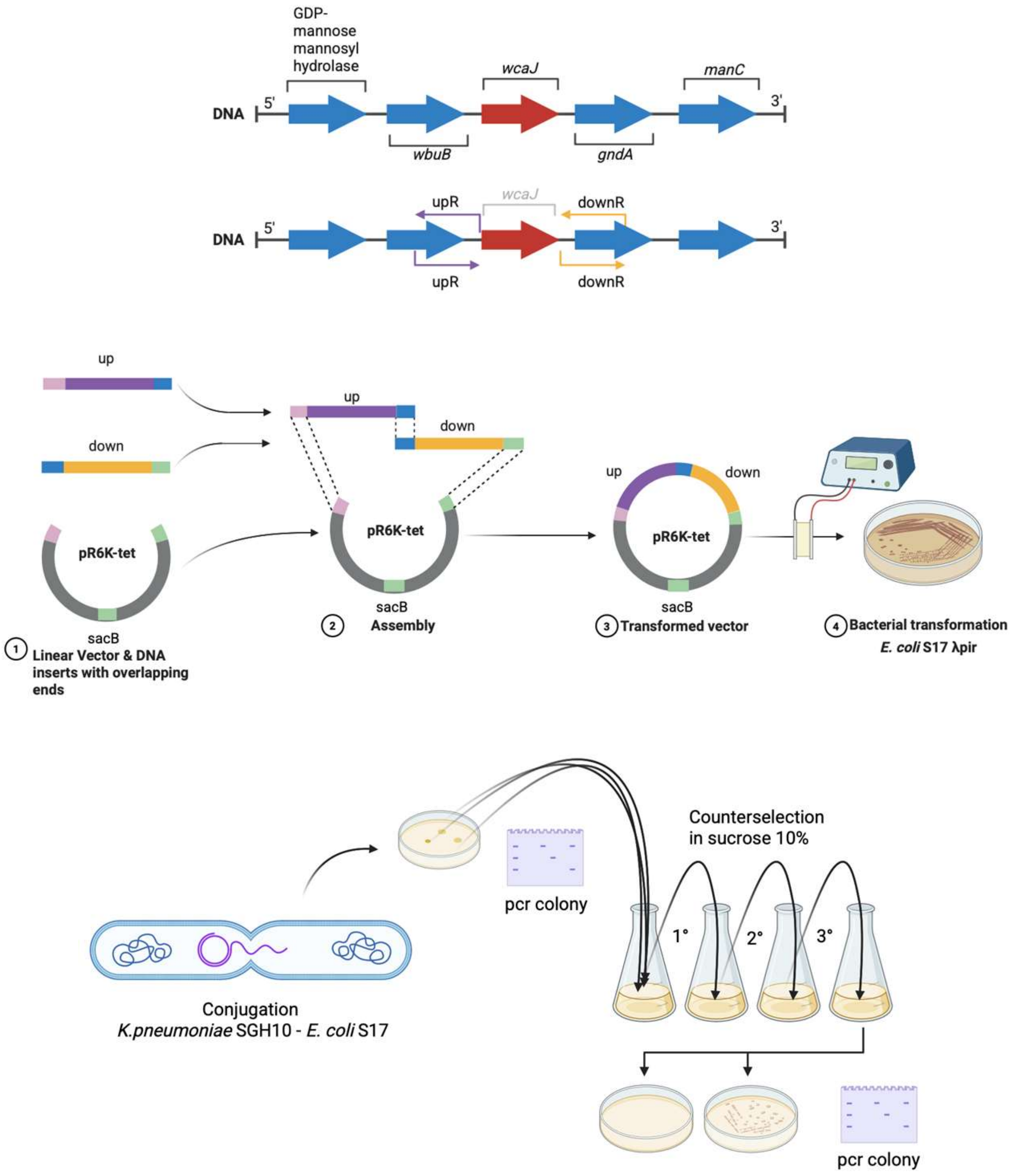


**Figure S1.** Overview of the site-directed mutagenesis strategy based on the suicidal plasmid pR6K-tet-sacB to generate the *K. pneumoniae* SGH10 Δ*wcaJ* deletion mutant. (A) Genetic environment of the *wcaJ* gene in the *K. pneumoniae* SGH10 chromosome. The purple and dark yellow arrows indicate the homology regions amplified by PCR to direct the precise deletion of the *wcaJ* gene. (B) Ligation of the upstream and downstream homology regions to the pR6K-tet vector using the NEBassembly kit and electroporation inyo the *E. coli* S17 λpir donor strain. (C) Negative selection based on the *sacB* gene and growth on 10% sucrose.

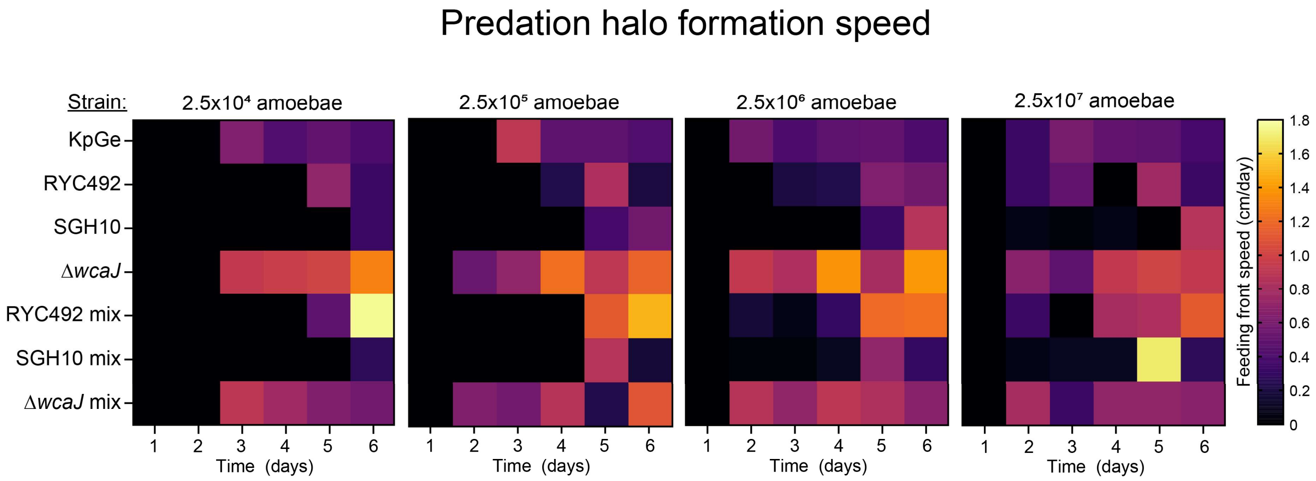


**Figure S2.** Predation halo formation speed of different amounts of *D. discoideum* cells after being seeded on a lawn of distinct *K. pneumoniae* strains. The feeding front speed was measured as centimeters per day.
